# Selective effects of cyclin dependent kinase inhibitors in gammaherpesvirus reactivation from latency

**DOI:** 10.64898/2026.03.18.712771

**Authors:** Joy E Gibson, Linda F van Dyk

## Abstract

Cell cycle manipulation is critical to oncogenesis, including cancers associated with oncogenic gammaherpesviruses, Epstein-Barr Virus and Kaposi’s Sarcoma-associated Herpesvirus. Infection with these viruses can result in various cancers, including lymphomas and carcinomas. In healthy individuals, gammaherpesvirus infections result in lifelong latent infections with occasional reactivation. The cell cycle plays a critical role in infection, particularly in reactivation from quiescent latency to lytic virus replication. A number of cyclin-dependent kinase (CDK) inhibitors are clinically available but with little investigation thus far for virus-associated cancers. Using the mouse gammaherpesvirus model, we assessed the impact of CDK inhibitors on virus reactivation. First, we tested chemical inducers of reactivation, and found that optimal reactivation occurred with a combination of PMA and sodium butyrate. Application of optimal reactivation triggers demonstrated distinct stage-specific outcomes of reactivation, distinguished using flow cytometry to measure expression of GFP (early reactivation) and vRCA, a late viral protein (late reactivation). Following chemical induction of reactivation, we used flow cytometry to demonstrate that the early effects of induction were unaffected by CDK inhibitors. However, all broad spectrum CDK inhibitors tested, Dinaciclib, Alvocidib, and Seliciclib, decreased both reactivation from latency and primary lytic replication. In contrast, the impact of targeted CDK 4/6 inhibitors, Palbociclib, Ribociclib, and Abemaciclib, was more nuanced, with decreased reactivation when given concurrently, but increased reactivation when administered prior to induction. These findings were consistent for both murine gammaherpesvirus and Epstein-Barr Virus. Overall, our data indicate that CDK inhibitors may be useful for targeted treatment of gammaherpesvirus-associated cancers, but optimal use of targeted CDK 4/6 inhibitors requires careful consideration of cell state and order of therapies.

## Introduction

Gammaherpesviruses are a subfamily of herpesviruses that have oncogenic potential, particularly in immune suppressed individuals. These include the human viruses Epstein-Barr Virus (EBV) and Kaposi’s Sarcoma-associated Herpesvirus (KSHV) and the mouse model, murine gammaherpesvirus 68 (γHV68). EBV infects more than 90% of the population worldwide^1^ and results in a range of pathogenic outcomes, including infectious mononucleosis, Burkitt’s lymphoma, Hodgkin’s lymphoma, nasopharyngeal carcinoma, and post-transplant lymphoproliferative disorders (PTLD).^2^ KSHV infection is less ubiquitous, yet results in significant morbidity from Kaposi’s sarcoma and primary effusion lymphoma.^2^ Despite the high prevalence of gammaherpesvirus infection and substantial associated morbidity and mortality, we lack effective therapies.

Gammaherpesvirus infection progresses through a well-defined sequence of events. Primary infection results in lytic infection with active replication and new virus production. This is followed by establishment of latency, predominantly in memory B cells.^3^ Latency persists for the life of the host and is characterized by viral transcriptional quiescence, with virus reactivation occurring spontaneously, or in response to a variety of triggers. Upon reactivation, latently infected cells initiate expression of viral genes and can re-enter a state of active viral replication, resulting in virus production and cell lysis.^4^ In both EBV and KSHV, latency genes are known to be critical for cellular transformation.^5, 6^ However, more recent studies have demonstrated that lytic genes are also important for oncogenesis.^6, 7^ Viral reactivation is typically rapidly controlled by the immune response, but in immune compromised states this can fail and may result in oncogenesis.^8–13^ Consideration of both reactivation and lytic infection is important for development of therapeutics.

KSHV and the mouse model gammaherpesvirus 68 (γHV68) encode a viral cyclin that mimics D-type cellular cyclins^14, 15^ and facilitates viral reactivation and pathogenesis.^16–21^ EBV instead regulates cellular D-type cyclins, which mediate oncogenesis.^22–24^ Cyclins are the regulatory binding partners for cyclin-dependent kinases (CDKs) and D-type cyclins bind to CDK4 and CDK6 to facilitate G1 cell cycle progression. Many studies demonstrate that CDKs cooperate with cyclins to support proliferation and influence the expression and activity of viral proteins.^25–28^ We propose that targeting this process through CDK inhibition may be a successful means to reduce viral reactivation and minimize pathogenic outcomes.

There are many CDK inhibitors being tested or approved for clinical use. Broad spectrum CDK inhibitors, including Dinaciclib, Alvocidib (Flavopiridol), and Seliclib (Roscovitine), have high toxicity but remain in clinical trials for a variety of cancers. Three selective CDK 4/6 inhibitors, Palbociclib, Abemaciclib, and Ribociclib, have been approved for clinical use for breast cancer. Ongoing clinical trials with these inhibitors are in progress for a variety of different cancers, including leukemia and lymphoma. No CDK inhibitor trials have specifically examined gammaherpesvirus-associated oncogenesis or treatment of gammaherpesvirus reactivation during immune compromised states. Early *in vitro* and murine studies using CDK inhibitors for treatment of gammaherpesvirus infection show promise. Alvocidib and Palbociclib suppress proliferation of EBV-infected lymphoblastoid cell lines.^26^ Recent studies identified Palbociclib as a potential therapeutic for EBV positive nasopharyngeal carcinoma in mice^29,30^ that can enhance KSHV tumor cell sensitivity to T cell killing.^31^ Palbociclib treatment of KSHV infected cells demonstrated a dependency on cellular cyclin^32^ and a metabolic requirement for the KSHV viral cyclin^33^. Abemaciclib was shown to be more potent than palbociclib in suppression of B-cell lymphoma cell proliferation and induction of apoptosis.^34^ Another broad spectrum CDK inhibitor, Alsterpaullone, resulted in decreased EBV positive tumor invasion and mortality.^35^ While early studies are promising, further comprehensive studies examining CDK inhibitors in EBV and KSHV oncogenesis are needed. Our work described here characterizes distinct stages of reactivation upon stimulation and assesses the impact of CDK inhibition on gammaherpesvirus reactivation using a murine lymphoma cell line, providing useful insights into the role of CDKs in reactivation and the potential for gammaherpesvirus treatment with CDK inhibitors.

## Materials and Methods

### Cell lines and reagents

A20-HE2.1 B cell lines (gift of Dr. Sam Speck, Emory University, Atlanta, GA) were cultured with 300 μg/mL Hygromycin B (Invitrogen) selection as described.^36^ Parental A20 B cells (ATCC TIB208) were cultured in RPMI 1640 (Gibco) supplemented with 10% bovine serum (FBS, Atlanta Biologicals) and 50 μM 2-mercaptoethanol (VWR). 3T12 mouse fibroblasts (ATCC CCL164) were cultured in DMEM supplemented with 5% FBS. MUTU I EBV-infected cells (gift from Dr. Shannon Kenney, University of WI, Madison) were cultured in RPMI 1640 with 10% FBS and used from passage 4-10. All cell culture media also contained 100 units/mL penicillin, 100 μg/mL streptomycin, and 2 mM _L_-glutamine (Gibco). Chemical inducers of reactivation: PMA (phorbol-12-myristate-13-acetate, 20 ng/mL), trichostatin A (5 μM), and bortezomib (20 nM) were obtained from Cell Signaling, and sodium butyrate (4 mM) from EMD Millipore. CDK inhibitors: Palbociclib (5 μM), Ribociclib (20 μM), Abemaciclib (1 μM), Dinaciclib (200 nM), Alvocidib (1 μM), and Seliclib (50 μM) were all obtained from Selleckchem and DMSO (Fisher Scientific) used as diluent. Infection of 3T12s was carried out with wild-type γHV68 clone WUMS (ATCC VR1465) at a multiplicity of infection of 5 pfu/cell.

### Quantitative PCR

For all quantitative PCR (qPCR) assays, cells were plated at a concentration of 5×10^5^ cells/mL (A20), 1×10^6^ cells/mL (MUTU I), or 1.5×10^5^ cells/mL (3T12) and treated for 24-72 hours. Cells were counted with trypan blue staining using an automated cell counter (Bio-Rad TC20 Automated Cell Counter). DNA was isolated using Qiagen DNeasy Kit per manufacturer protocol and quantified (NanoDrop ND-1000 Spectrophotometer) before suspension in DNase/RNase free water to 20 ng/μL. 25-100 ng was added to each qPCR reaction for final 20 ng/μL to allow comparison across experiments. Primers and probes: gB Forward 5’-GGCCCAAATTCAATTTGCCT-3’, gB Reverse 5’-CCCTGGACAACTCCTCAAGC-3’, and gB Probe 5’-FAM-ACAAGCTGACCACCAGCGTCAACAAC-TAMRA-3’, BALF5 Forward 5’-CGGAAGCCCTCTGGACTTC-3’, BALF5 Reverse 5’-CCCTGTTTATCCGATGGA ATG-3’, BALF5 Probe 5’-FAM-TGTACACGCACGAGAAATGCGCCT-BHQ-3’, NFAT5 Forward 5’-CATGAGCACCAGTTCCTACAATGAT-3’, NFAT5 Reverse 5’-TGCTTTGGATTTCGTTTTCGTGATT-3’, and NFAT5 Probe 5’ VIC-ACGAGGTACCTCAGTGTT-BHQ-3’. For BALF5 qPCR, results were normalized to cell number, calculated based on NFAT5 copy number (divided by 2 for NFAT5 per single cell). Standard curves were generated from plasmid standards. qPCR was completed with Roche LightCycler 480 Probe Master Mix on a QuantStudio 7 Flex thermocycler (Applied Biosystems) with the settings: 95°C for 3 minutes followed by 40 cycles of 95°C 10 seconds 55°C 30 seconds. Copy number per reaction was calculated using the QuantStudio Real-Time PCR Software based on standard curve.

### Flow cytometry

Cells were stained with LIVE/DEAD Fixable Near-IR Dead Cell Stain Kit (Invitrogen) per manufacturer protocol. Cells were fixed with 2% paraformaldehyde (Alfa Aesar). Cells were incubated with 1:5000 dilution of rabbit anti-vRCA antibody (gift of Dr. H.W. Virgin, Washington University), conjugated to PE by Zenon labeling kit (Invitrogen), and purified 2.4G2 antibody (Tonbo). LSR Fortessa X-20 (BD) data was analyzed using FlowJo software (BD). Cells were gated for doublet and dead cell exclusion prior to GFP and vRCA-PE analysis, with a single exception noted in Figure 3, and compensation calculated and corrected visually based on single stain controls. To account for GFP expression in A20-HE2.1 cells, single stain controls included the GFP negative parental A20 cells and lytically infected 3T12 cells.

### Statistical analyses and calculations

All results were analyzed using paired T-test calculated with GraphPad Prism, with Grubbs’ test for outliers applied to all. Cell doubling time was assessed using the following equation: Doubling time = ((duration*log(2))/(log(FinalConcentration)-log(InitialConcentration))).

## Results

### Reactivation in the A20-HE2.1 cell line is optimally induced by PMA and NaB

The experiments described herein use a B cell lymphoma cell line, A20-HE2.1,^36^ that is latently infected with γHV68. These cells harbor a full γHV68 genome with a viral transgene containing an expression cassette encoding hygromycin resistance fused to GFP under control of the human cytomegalovirus promoter (Figure 1A). When treated with chemical stimuli, A20-HE2.1 cells can initiate lytic gene expression through the process of reactivation. Reactivation leads to virion production and release. We used GFP expression, viral DNA amplification, and cell surface expression of late viral proteins as readouts of reactivation.

**Figure 1.**
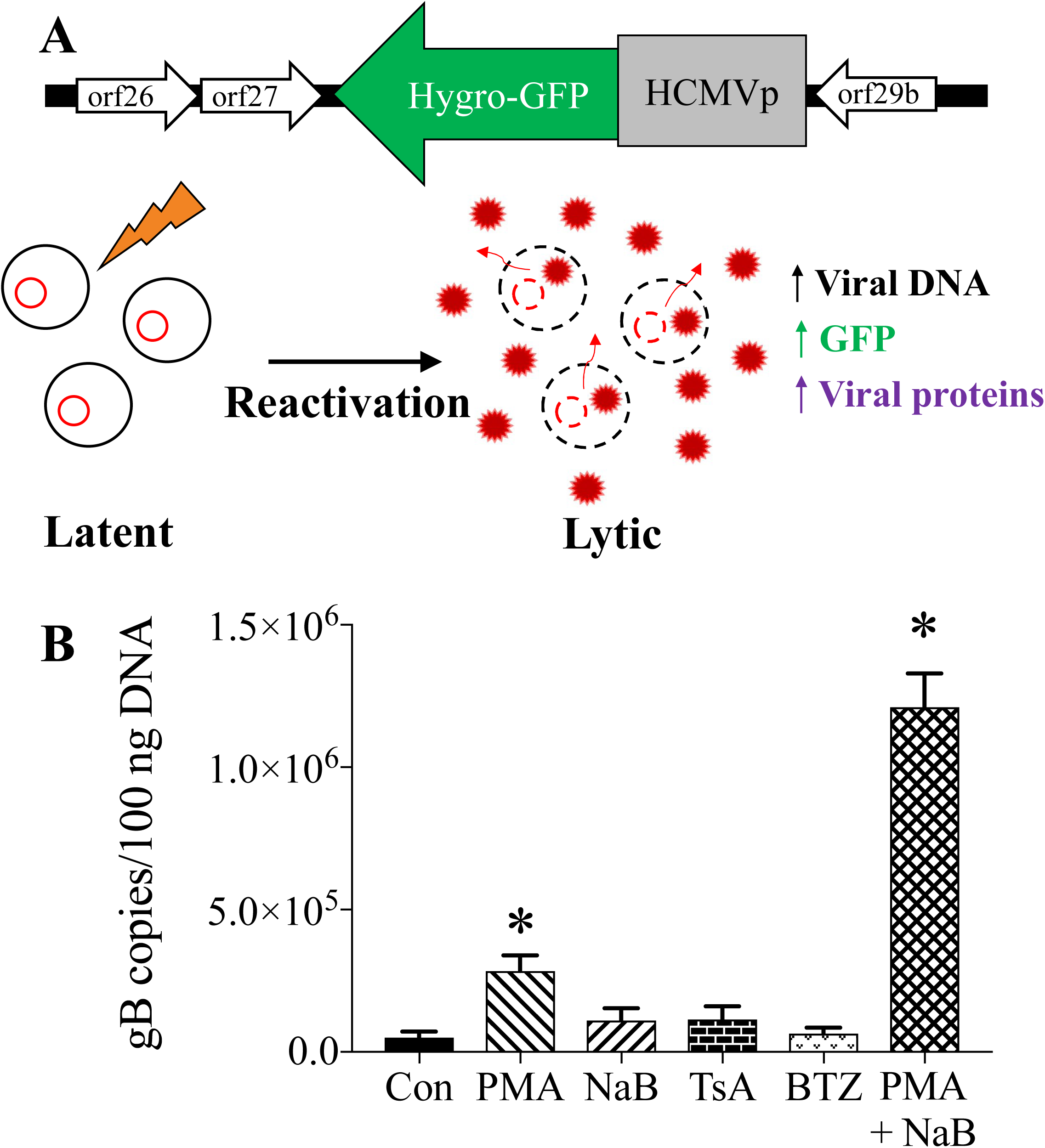
Reactivation in the A20-HE2.1 cell line is optimally induced by combination treatment with PMA and NaB. Diagram of murine A20-HE2.1 cells **(A)** with a γHV68 genome that contains a hygromycin cassette fused to GFP, expressed under a human CMV promoter. These cells can be induced to reactivate from latency (left) to lytic infection (right), measured by an increase in viral DNA, GFP expression, and viral proteins. **(B)** Viral replication quantified by qPCR gB copies in A20-HE2.1 cells treated for 24 hours with PMA, NaB, TsA, BTZ, or both PMA and NaB (PMA+NaB). Controls (Con) were treated with vehicle (ethanol or DMSO), shown as mean +/-SEM. (*p<0.05, n=6 Con, PMA, NaB, n=4 TsA, BTZ, n=3 PMA+NaB)

To determine the optimal inducer(s) of reactivation, we tested four known reactivation inducers, PMA (a protein kinase C activator), sodium butyrate (NaB, a histone deacetylase inhibitor), trichostatin A (TsA, a histone deacetylase inhibitor), and bortezomib (BTZ, a proteasome inhibitor). Combination treatment with PMA+NaB led to optimal reactivation, with a 40-fold increase in viral DNA content from 29,041 copies to 1,152,808 copies per 100 ng DNA, determined by qPCR (Figure 1B). Treatment with PMA alone also significantly increased reactivation, with an 11-fold increase in viral genome copy number to 311,758 copies per 100 ng DNA (Figure 1B). NaB, TsA, and BTZ did not significantly increase viral DNA production.

### Reactivation stimulation results in differential induction of viral gene expression

We used flow cytometry to evaluate viral gene expression during chemical reactivation (gating strategy shown in Figure 2A). GFP transgene expression from the virus genome served as a marker of genome accessibility and viral gene expression. Viral regulator of complement activity (vRCA) protein expression was a marker of late reactivation, as this protein is produced late during reactivation and present on the cell and virion envelopes. PMA+NaB treatment led to increased GFP expression from 7.3% (+/-1.2) to 75.1% (+/-6.6) and vRCA from 0.2% (+/-0.08) to 3.2% (+/-0.4), with far more GFP positive than vRCA positive cells (Figure 2 B/C). Mean fluorescence intensity (MFI) of GFP correlated with these data and was higher with PMA+NaB treatment. In cells treated with reactivation inducers, most GFP positive cells remain vRCA-PE negative (97%+/-0.4), while vRCA-PE positive cells are also GFP positive (89% +/-5.2) (Figure 2D).

**Figure 2.**
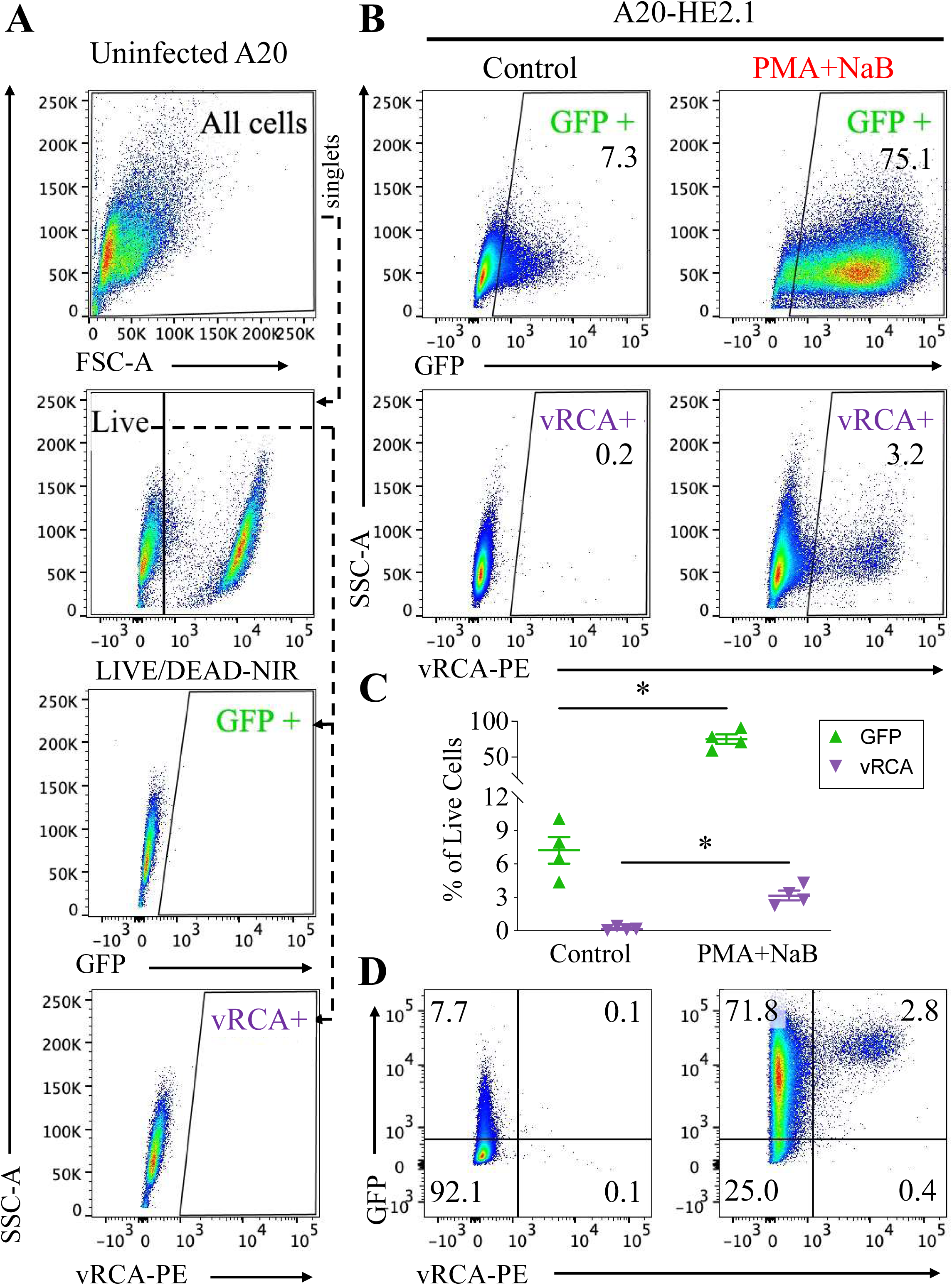
Chemical induction of reactivation increases viral gene expression in most cells, but only a small percentage of cells progress to lytic infection. Flow cytometry of A20-HE2.1 cells stained with PE-tagged antibody against vRCA (vRCA-PE) and a LIVE-DEAD stain with Near IR (NIR) fluorescence (LIVE/DEAD-NIR). **(A)** Representative flow cytometry plots demonstrating gating strategy: all cells -> side scatter singlets -> forward scatter singlets -> live cells->GFP or vRCA. Live cells identified by negative NIR fluorescence. **(B,C)** Mean +/- SEM of GFP and vRCA expression in control-treated (left) and 24 hour PMA+NaB-treated (right) cells with representative flow cytometry scatter plots. **(D)** Representative flow cytometry biaxial plots with mean expression in each quadrant for GFP vs. vRCA expression. (*p<0.05, n=4)

GFP and vRCA expression were compared for different chemical inducers of reactivation (Figure 3A/B). With treatment, GFP increased from 3.3% (+/-1.9) - 6.1% (+/-1.1) of cells (EtOH, DMSO) to 35.3% (+/-10.4) (PMA), 17.5% (+/-4.4) (NaB), 60.8% (+/- 16.5) (TsA), 73.6% (+/-8.4) (BTZ), or 68.9 % (+/-2.9) (PMA+NaB). PMA+NaB was the only treatment that significantly increased vRCA expression, from 0.2% (+/-0.07) to 2.3% (+/-0.5). These treatments all resulted in ready initiation (GFP) of viral gene expression with limited progression to late (vRCA) gene expression.

**Figure 3.**
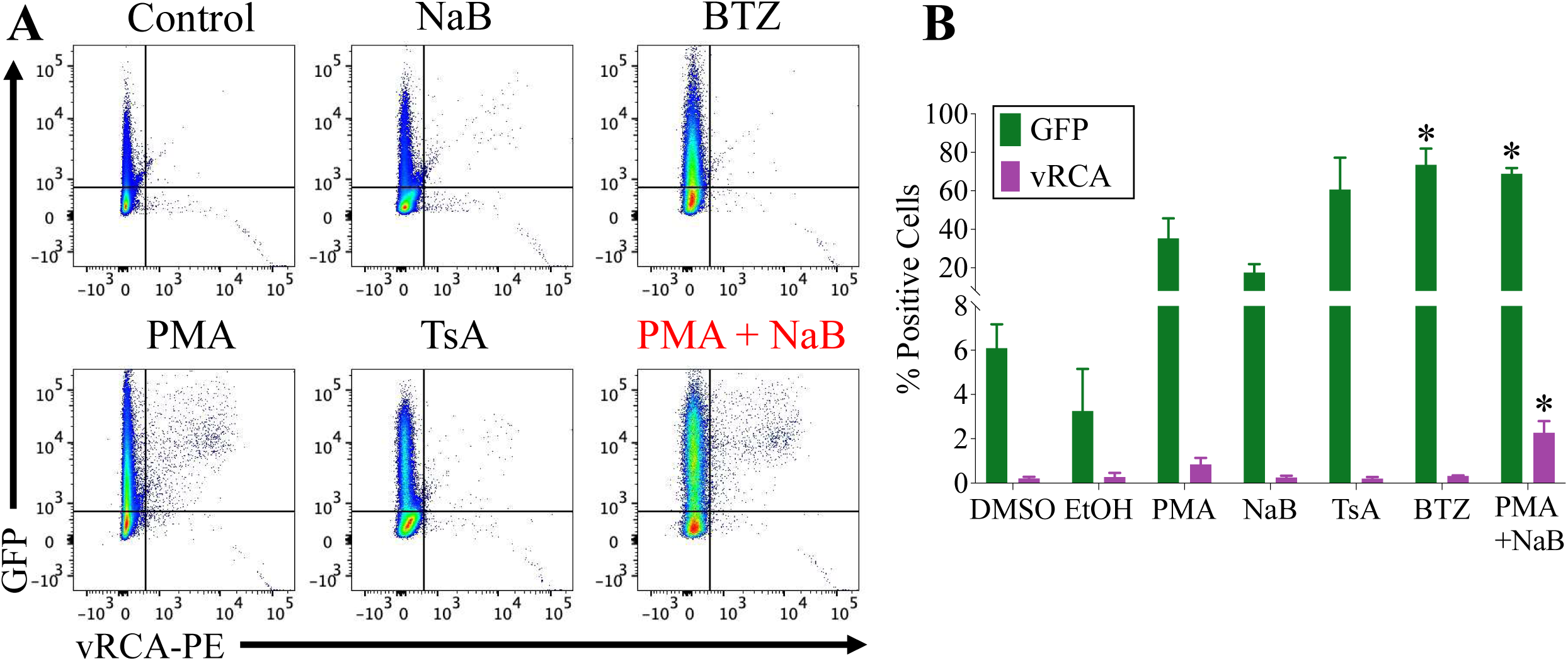
Weak chemical inducers of reactivation upregulate viral gene expression without progression to lytic infection. A20-HE2.1 cells treated with vehicle (DMSO or ethanol, EtOH), PMA, NaB, TsA, BTZ, or PMA+NaB for 24 hours. **(A,B)** Mean +/-SEM of GFP and vRCA expression by flow cytometry following chemical treatment, with representative biaxial plots showing GFP expression (y-axis) and vRCA-PE expression (x-axis). (*p<0.05, n=3 except PMA+NaB, n=8)

### Clinical CDK inhibitors impact viral reactivation and lytic infection in a time sensitive manner

We examined the impact of the broad spectrum CDK inhibitor Dinaciclib (CDK 1, 2, 5, 9 inhibitor) and the specific CDK 4/6 inhibitor Palbociclib on reactivation. In A20-HE2.1 cells, concurrent treatment with reactivation inducers resulted in decreased viral reactivation, as measured by viral DNA amplification, 0.68-fold with Palbociclib and 0.04-fold with Dinaciclib (Figure 4A). The response to Palbociclib, but not Dinaciclib, was dependent on treatment timing. Pre-treatment with Palbociclib resulted in a 2.52-fold increase in reactivation, while Dinaciclib pre-treatment potently inhibited reactivation 0.03-fold (Figure 4B).

**Figure 4.**
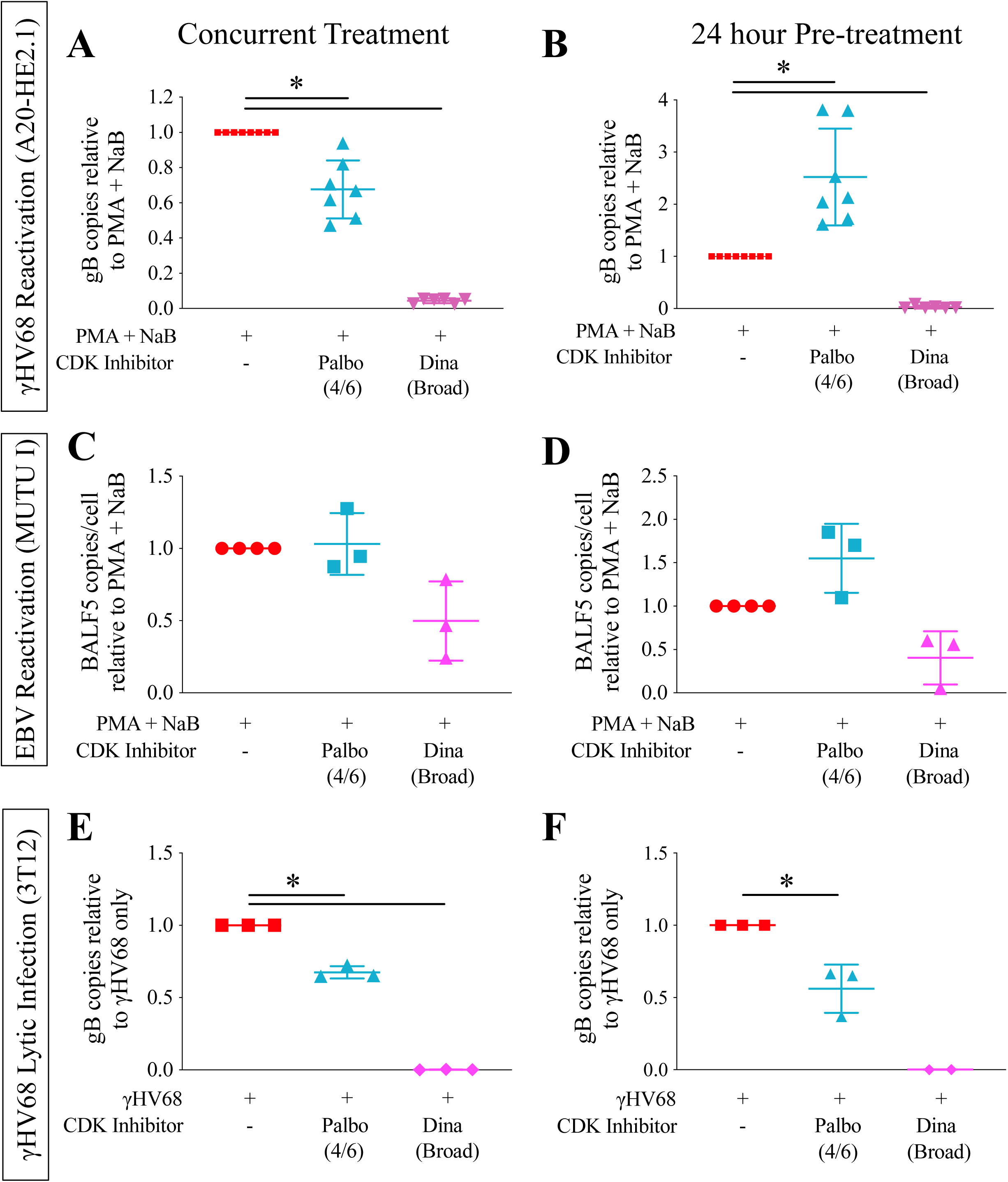
CDK inhibition impacts viral reactivation and lytic infection in a time sensitive manner. Cells were treated with vehicle (water and DMSO), Palbociclib (Palbo), or Dinaciclib (Dina) either concurrently **(A, C, E)** or 24 hours prior to **(B, D, F)** reactivation induction or infection. Reactivation in A20-HE2.1 cells **(A, B)** or EBV positive MUTU I cells **(C, D)** was induced by treatment with PMA+NaB for 24 hours (A20-HE2.1) or 48 hours (MUTU I). Reactivation control was treated with vehicle only (water and DMSO). A permissive murine fibroblast cell line (3T12) was lytically infected with γHV68 at an MOI of 5 for 24 hours **(E, F)**. Quantification of viral copies was done using qPCR for the viral gene gB (γHV68) or BALF5 (EBV). For MUTU I cells, BALF5 copy number was normalized to cell number using qPCR against host cell NFAT5. Both gB and BALF5 were normalized to PMA+NaB alone to give relative quantity. (*p<0.05, n=7 for A and B, n=3 for C-F)

We also examined the impact of these inhibitors on reactivation of EBV. Using a Burkitt lymphoma cell line latently infected with EBV, reactivation can be induced by treatment with PMA+NaB for 48 hours. The response of EBV to CDK inhibition was more variable but pre-treatment with Palbociclib increased viral DNA copies 1.55-fold, while concurrent treatment had no effect (Figure 4C/D). Dinaciclib reduced viral DNA copy number 0.4- to 0.5-fold regardless of treatment timing, less potently than γHV68 reactivation (Figure 4C/D).

Viral D-type cyclins are not known to play a role in primary lytic infection. Therefore, we also examined the impact of CDK inhibition on primary lytic infection of mouse 3T12 fibroblasts. Both Dinaciclib and Palbociclib inhibited lytic infection regardless of treatment timing, 0.002-fold and 0.6- to 0.7-fold, respectively (Figure 4E/F).

We then tested related CDK inhibitors to determine if the effects were consistent across inhibitor types. In addition to Dinaciclib, we tested the broad spectrum CDK inhibitors Alvocidib (flavopiridol; CDK 1, 2, 4, 6, 9 inhibitor) and Seliciclib (roscovitine; CDK 2, 5, 7, 9 inhibitor). Broad spectrum CDK inhibitors resulted in strong inhibition of γHV68 reactivation 0.02- to 0.05-fold regardless of the inhibitor or treatment timing (Figure 5B).

**Figure 5.**
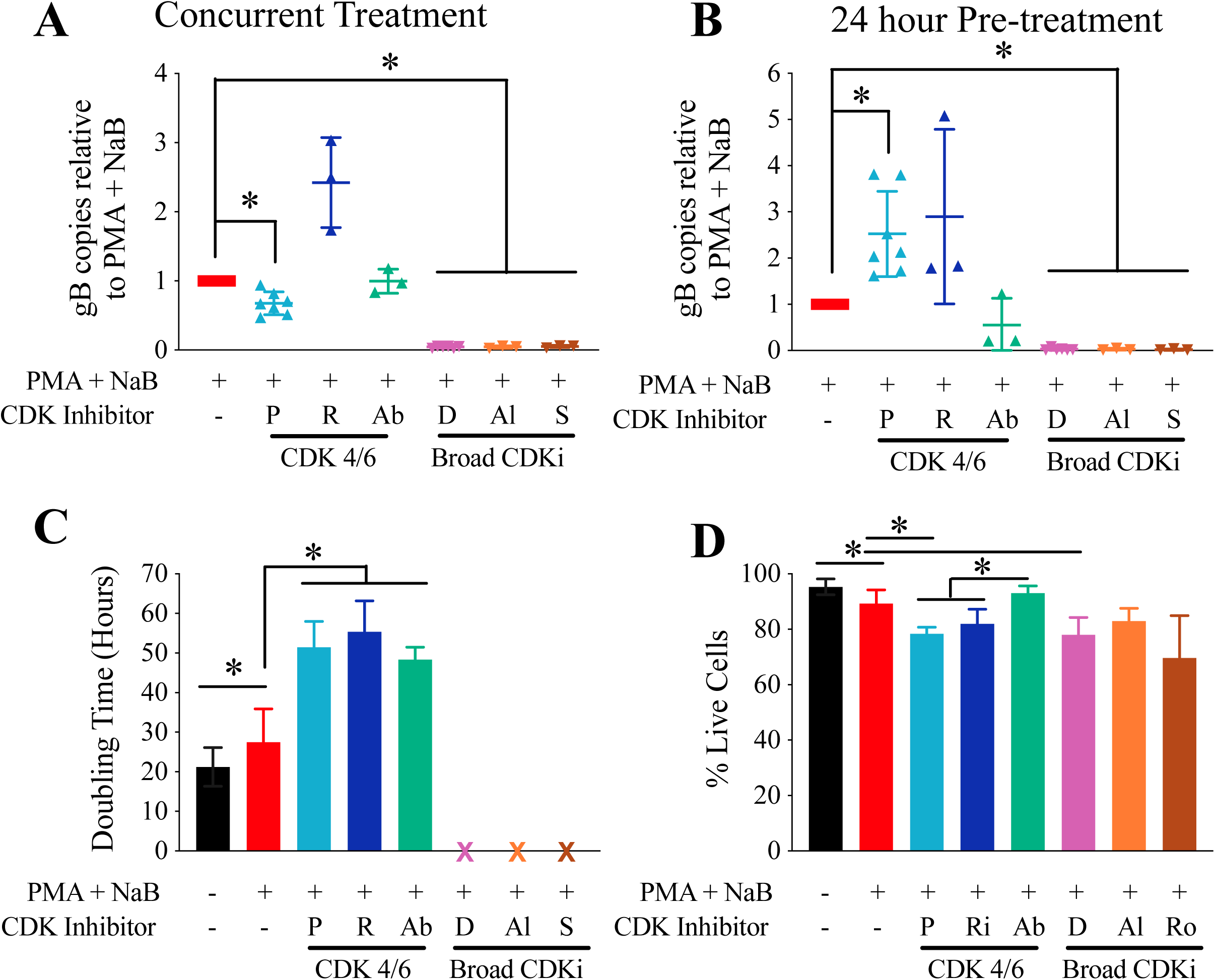
Effect of CDK 4/6 inhibition varies with inhibitor. A20-HE2.1 cells were treated with vehicle (water and DMSO), Palbociclib (P), Ribociclib (R), Abemaciclib (Ab), Dinaciclib (D), Alvocidib (Al), or Seliciclib (S). Reactivation in murine A20-HE2.1 cells was induced by treatment with PMA+NaB for 24 hours. **(A,B)** qPCR for the viral gene gB, represented by mean +/-SEM gB copies relative to PMA+NaB alone during concurrent **(A)** and pre-treatment **(B)** with inhibitor. **(C)** A20-HE2.1 cell doubling time with PMA+NaB treatment and CDK inhibitors, based on trypan blue counting. **(D)** Viability (percentage of cells that are alive) in cells treated with PMA+NaB and CDK inhibitors, mean +/-SEM. (*p<0.05; A and B n=8 for Palbociclib and Dinaciclib, n=3 for all other inhibitors; C and D n=6 for Control, PMA + NaB, Palbociclib, Dinaciclib, n=3 for all other inhibitors)

In addition to Palbociclib, we tested the specific CDK 4/6 inhibitors Ribociclib and Abemaciclib. Effects of these inhibitors on γHV68 reactivation in A20-HE2.1 cells were more variable than that of Palbociclib. In contrast to a decrease with Palbociclib, concurrent treatment with Ribociclib increased viral DNA 2.5-fold but Abemaciclib had no effect (Figure 5A). With pre-treatment, both Palbociclib and Ribociclib increased viral DNA 2.5- and 2.9-fold, respectively, while Abemaciclib decreased viral DNA 0.6-fold (Figure 5B).

Variation in the effects of different CDK inhibitors did not appear to be due to different effects on cell cycle progression or viability. Cell doubling time was increased from 27.5 hours to 48-55 hours with all CDK4/6 inhibitors, consistent with inhibition of cell division. Of note, rate of cell division did slow with PMA+NaB treatment alone, resulting in an increased doubling time to 27.5 hours from 21.2 hours (Figure 5C). The broad spectrum CDK inhibitors all completely halted cell division. Regarding viability, treatment with PMA+NaB decreased viability by 6% (Figure 5D). All CDK inhibitors further decreased cell viability by about 10%, except for Abemaciclib, which resulted in slightly less cell death than treatment with PMA+NaB alone (Figure 5D).

### CDK inhibition alters progression to late stage reactivation

We used flow cytometry to assess the impact of CDK inhibition on early and late stage reactivation by comparing GFP and vRCA expression. Reactivation induction resulted in increased GFP expression in all conditions regardless of type or timing of CDK inhibition (Figure 6). However, while still enhanced from baseline, GFP expression was decreased with Palbociclib treatment regardless of whether cells were treated concurrently (80.1% +/-6.4 to 58.1% +/-8.6) or pre-treated (77.3% +/-3.3 to 69.2% +/-2.7). Conversely, changes in vRCA late gene expression correlated with changes in viral DNA copy number. In cells treated concurrently, vRCA expression decreased with Dinaciclib treatment and was unchanged with Palbociclib treatment (Figure 6B). In cells pre-treated with CDK inhibitors, vRCA expression significantly decreased with Dinaciclib treatment, and increased 4-fold with Palbociclib treatment, consistent with DNA replication findings (Figure 6C).

**Figure 6.**
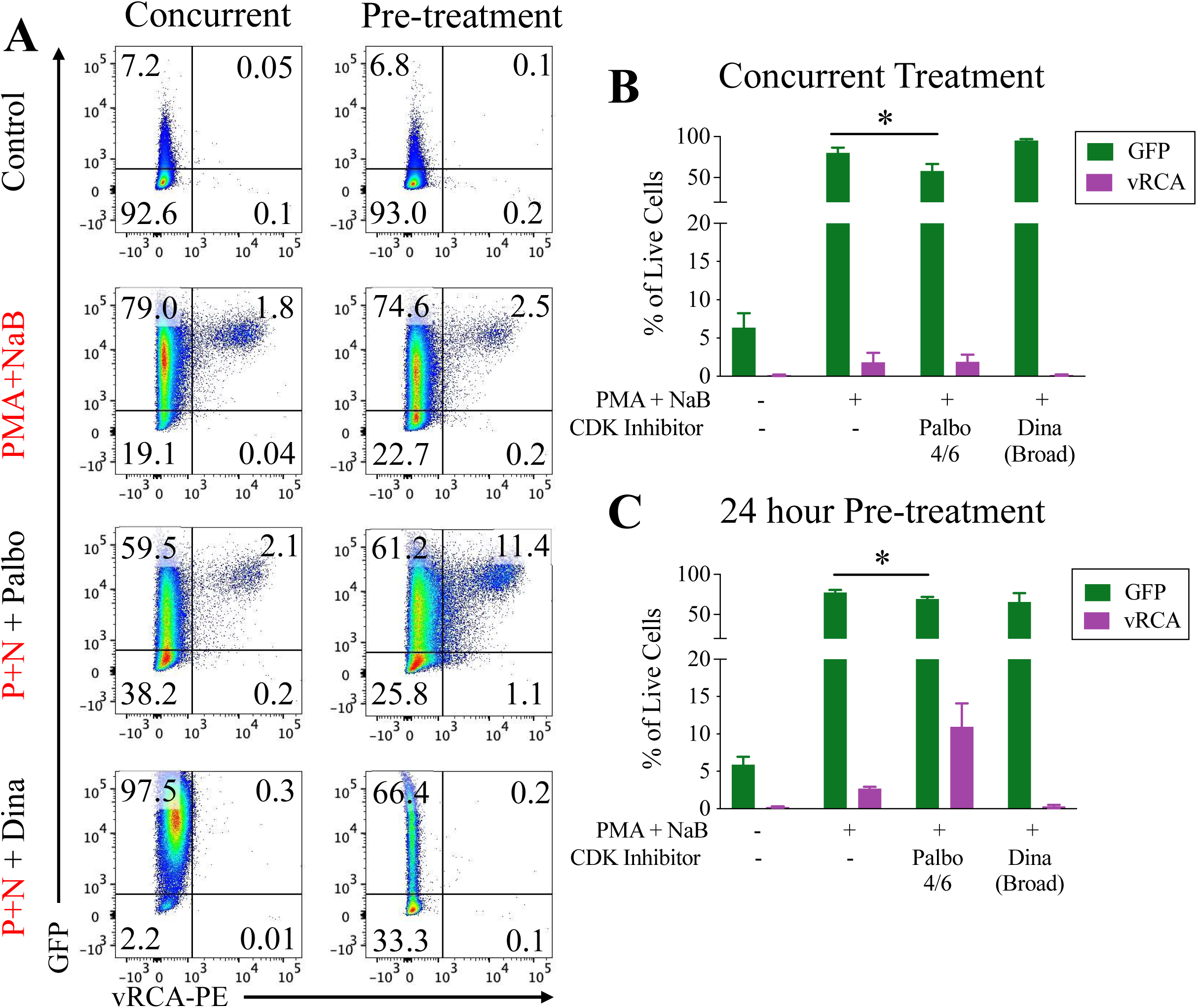
CDK inhibition alters viral gene expression and progression to lytic virus production. A20-HE2.1 were treated with vehicle (water or DMSO), Palbociclib (Palbo), and Dinaciclib (Dina) either concurrent or 24 hours prior to reactivation induction. **(A)** Representative flow cytometry plots showing cells treated concurrently (left) or 24-hours prior to treatment (right). After gating to exclude doublets and dead cells, GFP expression (y-axis) and vRCA-PE expression (x-axis) were analyzed. Mean +/-SEM of expression in cells treated with PMA+NaB and CDK inhibitors concurrently **(B)** or 24 hour prior to PMA+NaB **(C)**. (*p<0.05, n=3)

### Differential response to CDK inhibitors does not appear to be related to toxicity

To determine whether some CDK inhibitors caused more cellular toxicity than others, we assessed cellular viability using two methods. First, we used trypan blue exclusion to determine A20-HE2.1 cell viability. Dinaciclib decreases cell viability in both control and PMA+NaB-treated cells (Figure 7A). However, Palbociclib significantly decreased viability only in PMA+NaB-treated cells (Figure 7A). Pre-treatment with CDK inhibitors yielded similar results (data not shown).

**Figure 7.**
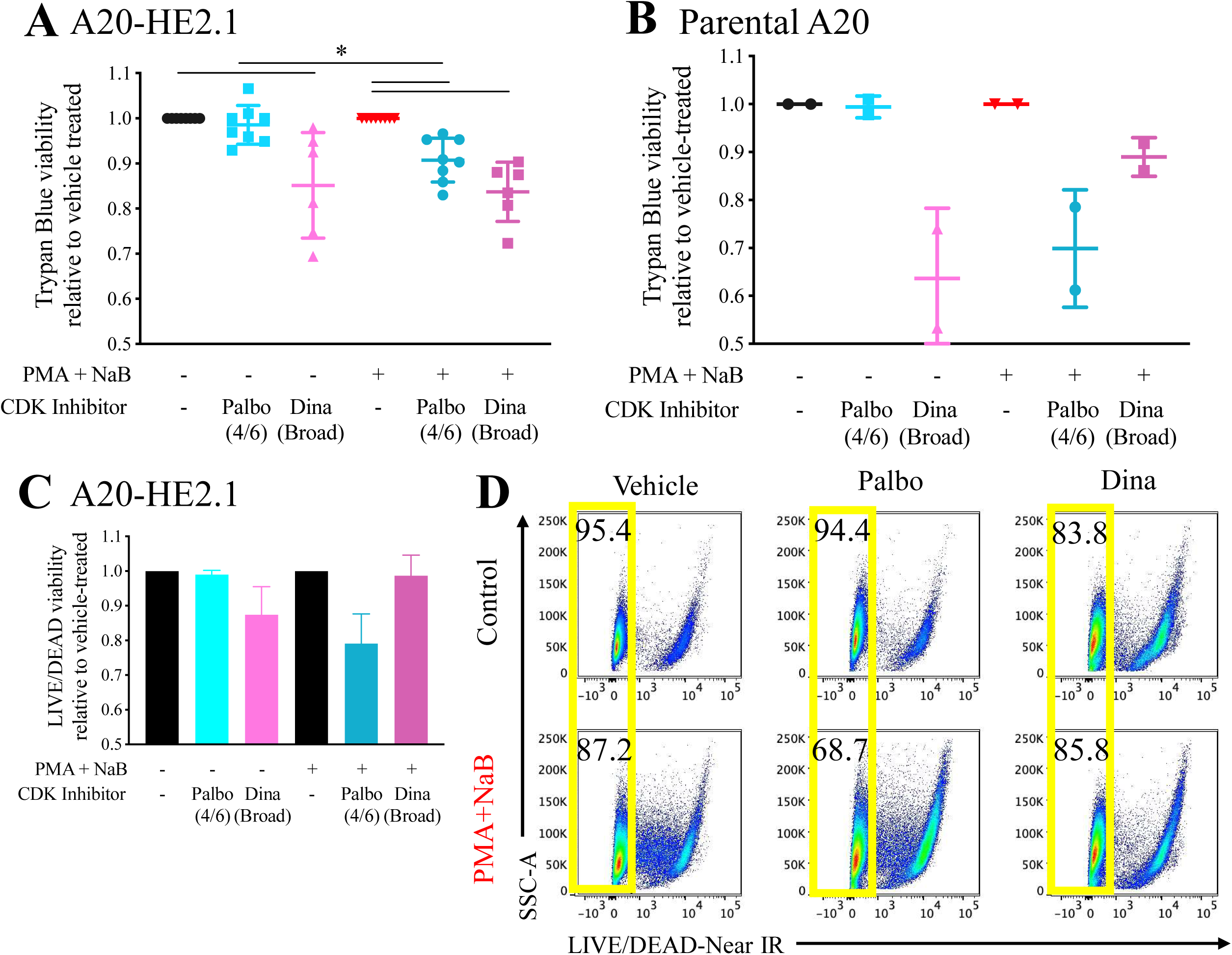
CDK4/6 inhibition selectively reduces cell viability in PMA+NaB-treated cells. A20-HE2.1 or uninfected (parental) A20 cells were treated with vehicle (water or DMSO), Palbociclib (Palbo), or Dinaciclib (Dina) concurrent with PMA+NaB for 24 hours. Cell viability was quantified by trypan blue staining or flow cytometry using LIVE/DEAD-Near IR stain. Viability was normalized to vehicle-treated condition for control or PMA+NaB. Mean relative viability +/-SEM based on trypan blue staining shown for A20-HE2.1 **(A)** or parental A20 **(B)** cells. Mean relative viability +/- SEM based on LIVE-DEAD-Near IR staining **(C)**, with representative flow cytometry plots showing percentage of cells that are live (yellow) **(D)**. (*p<0.05; A: n=8 for Palbo, n=6 for Dina; B: n=2; C: n=3)

The selective toxicity of Palbociclib could be due to viral reactivation or the combined chemical treatment. To address this, we quantified cell viability in uninfected parental A20 cells and found that Palbociclib and Dinaciclib followed a similar pattern. Dinaciclib decreased relative viability in both control and PMA+NaB-treated cells, while Palbociclib only decreased relative viability in PMA+NaB-treated cells (Figure 7B). Trypan blue results were confirmed using flow cytometry with a LIVE/DEAD stain to quantify cell viability in PMA+NaB and CDK inhibitor treated cells. Although the overall trend appeared similar, viability was more variable using this technique. Dinaciclib decreased relative viability in control but not PMA+NaB-treated cells, while Palbociclib decreased relative viability in PMA+NaB-treated cells but not control cells (Figure 7C/D).

## Discussion

In this paper, we presented two major findings. First, we identified that viral reactivation is a measurably stepwise process, with distinct early and late stages of reactivation quantifiable by flow cytometric analysis. Second, we characterized the effect of CDK inhibitors on gammaherpesvirus latency and reactivation toward their potential to treat gammaherpesvirus-associated cancers. We found that broad spectrum CDK inhibitors universally inhibit both lytic virus infection and viral reactivation, while the response to CDK 4/6 inhibition is more complex and has potential to more specifically target latency and reactivation.

To distinguish early and late stage reactivation, we used GFP and vRCA (a late lytic protein) expression in the A20-HE2.1 cell line to separate three distinct populations: latent (GFP-vRCA-), early reactivation (GFP+ vRCA-), and late reactivation (GFP+ vRCA+). This distinction allowed us to identify unique steps in the response to chemical reactivation induction. Interestingly, all of the chemical inducers of reactivation were competent to induce early reactivation (initiation of reactivation), while only PMA+NaB and, to a lesser extent, PMA alone, supported progression to late stage reactivation. PMA is a protein kinase C activator that has been shown to induce MAPK/Erk signaling pathways^37^ as well as cell cycle arrest^38^ in reactivating cells. Therefore, it is likely these processes are important for viral reactivation. The synergy between sodium butyrate and PMA seen here suggests that sodium butyrate induction of early reactivation may increase the ability of PMA to drive progression to late reactivation. As trichostatin A and bortezomib appeared to increase early reactivation to a greater extent than sodium butyrate, it is possible that combination treatments may lead to more potent reactivation induction that could be optimized for different infected cell types and states. These findings provide useful insight into the mechanism of reactivation and may allow more targeted reactivation induction in future studies.

CDK inhibitors show significant promise clinically for cancer treatment, particularly the less toxic targeted CDK 4/6 inhibitors.^39, 40^ The use of these inhibitors for treatment of oncogenic viral infections has not been thoroughly explored. Despite their unique origin and pathogenesis, virus-associated cancers are generally treated similarly to other cancers. Antivirals that are effective for treatment of other herpesviruses, such as acyclovir, ganciclovir, and cidofovir, are not effective clinically against EBV or KSHV-associated cancers.^41, 42^ This has been attributed to the fact that virus within tumors is in a state of latency and, therefore, resistant to antivirals that inhibit viral DNA synthesis.^43^ Even in lytic infection states such as infectious mononucleosis or EBV viremia following transplantation, the utility of antiviral therapy is controversial.^44–47^ CDK inhibitors may provide a new approach to antiviral treatment targeting both viral and cellular processes in reactivation and oncogenesis and may provide a useful new component of combination therapies.

We demonstrated that the broad spectrum CDK inhibitors Dinaciclib, Alvocidib, and Seliciclib consistently inhibit viral reactivation of γHV68 and EBV, and lytic infection with γHV68. CDK inhibition seems to target the progression from early to late stage reactivation (model shown in Figure 8). Therefore, it is likely CDK4/6 activity is required for late steps in reactivation and the viral lytic cycle. Alvocidib and Seliciclib have been shown to inhibit other herpesviruses^48–50^ and have shown promise for treatment of non-herpes viruses.^51–55^ Our data demonstrate they may also be effective for treatment of gammaherpesvirus infection.

**Figure 8.**
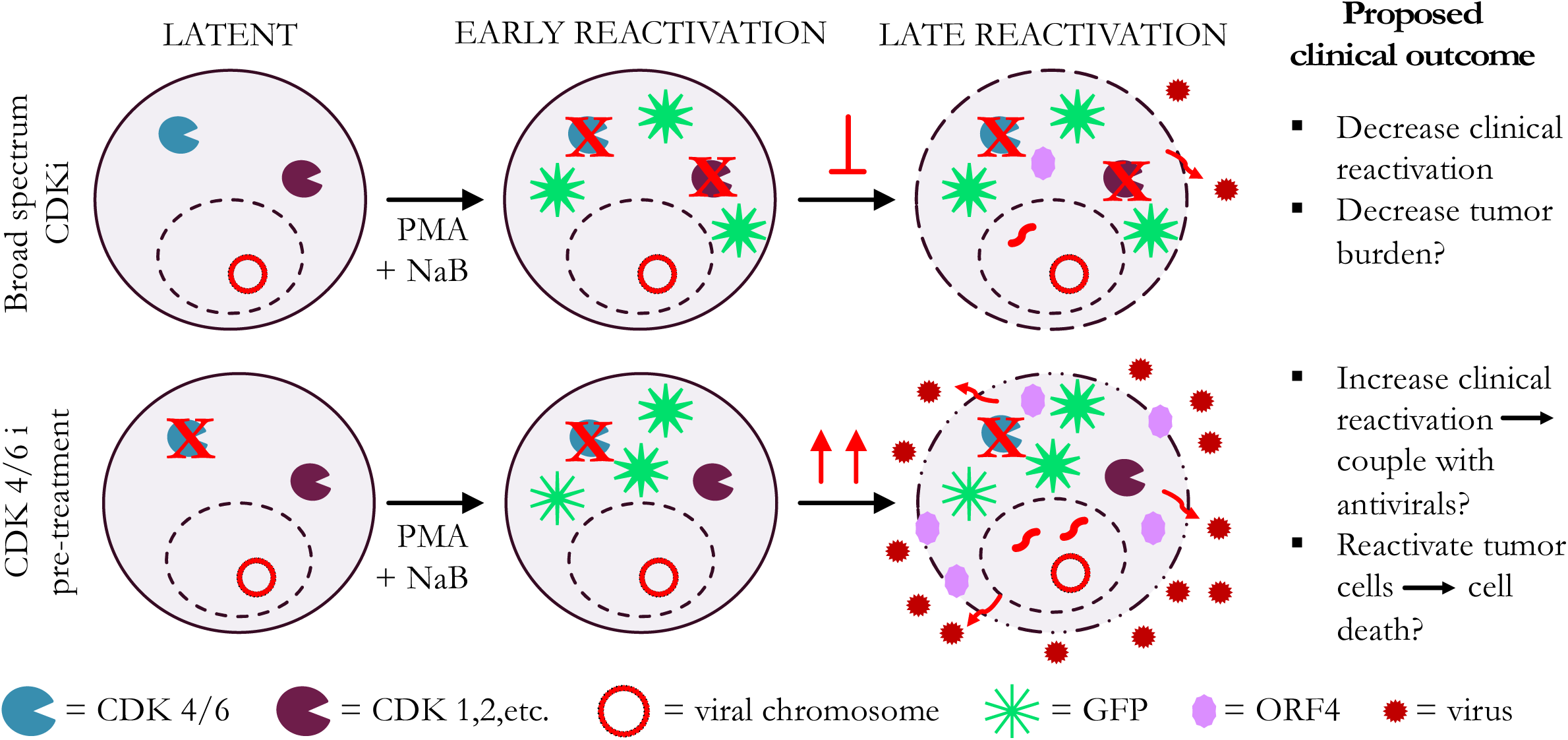
Working model for the role of CDK inhibitors in gammaherpesvirus reactivation. During PMA+NaB treatment with broad spectrum CDK inhibitors, early viral reactivation occurs, as demonstrated by increased GFP expression. However, there is potent inhibition of the progression to late stages of viral reactivation, as shown by decreased vRCA expression and decreased viral DNA copies, a marker of viral replication. This suggests that broad spectrum CDK inhibitors may be useful to decrease viral reactivation in immune compromised patients. It may also have the potential to decrease gammaherpesvirus-associated tumor burden. On the contrary, pre-treatment of cells with CDK 4/6 inhibitors, particularly Palbociclib and Ribociclib, enhances viral reactivation. When CDK 4/6 is inhibited while cells are still in latency, early reactivation, as demonstrated by GFP expression, seems to continue. These cells seem to be more susceptible to reactivation, with increased late reactivation as demonstrated by increased vRCA expression and increased viral DNA copies.

For the targeted CDK 4/6 inhibitors Palbociclib, Ribociclib, and Abemaciclib, our findings were more nuanced and specific to distinct stages in virus reactivation. Concurrent treatment with CDK 4/6 inhibitors and reactivation inducers had a similar effect to broad spectrum CDK inhibitors. However, 24-hour pre-treatment generally increased late stage reactivation in γHV68 and EBV, with increased viral DNA and vRCA. Interestingly, initiation (early stage, GFP expression) of reactivation was not increased by CDK 4/6 inhibitor pre-treatment, which further supports the conclusion that CDK 4/6 activity drives progression from early to late stage reactivation. The enhanced progression to late reactivation that occurs with CDK 4/6 inhibition suggests that CDK 4/6 inhibition results in a cellular state that is more favorable for completion of reactivation. This could be due to a number of potential factors. One possibility is that cells in the G_0_/G_1_ phase of the cell cycle are more susceptible to reactivation. Some studies have suggested that gammaherpesvirus reactivation results in cell cycle arrest at G_0_/G_1_,^56, 57^ likely related to a global host shutoff that allows viral gene expression to predominate. A number of reactivation inducers are known to cause cell cycle arrest in infected cells.^38, 58, 59^ However, CDK 4/6 inhibition alone did not induce reactivation (data not shown) and the inhibition of reactivation when given concurrently suggests that the mechanism of action is more complex.

CDK 4/6 inhibition may increase reactivation capability in other ways. The CDK 4/6 inhibitors used are competitive inhibitors, which would likely result in increased free cellular viral cyclin or D-type cyclins. This may lead to CDK-independent, or CDK4/6-independent, effects of cyclins that have the potential to promote reactivation. D-type cyclins have a number of CDK-independent effects mediated through transcription regulation or direct protein binding that regulate cell function, including cell proliferation and migration, cell metabolism, p53-mediated cell cycle control, DNA repair, and programmed cell death.^60–64^ It is possible that these CDK-independent functions of D-type cyclins are increased with CDK 4/6 inhibition and enhance viral reactivation. Conversely, CDK6 has been identified to have several cyclin and kinase-independent functions that are not complemented by CDK4, including hematopoietic stem cell differentiation and leukemia formation through Egr1 regulation.^65^ Egr1 is known to play a key role in EBV lytic infection and reactivation and induces cyclin D1 expression.^66–72^ Therefore, it is possible that CDK6 could increase Egr1 expression to support reactivation. Competitive CDK4/6 inhibition may free up CDK6 for kinase-independent functions and drive reactivation, as seen in our model.

Our findings that pre-treatment with CDK4/6 inhibitors results in enhanced viral reactivation yield several new insights into potential mechanism for viral reactivation and for the clinical potential of these inhibitors, which warrant further investigation. In the setting of clinical reactivation following transplantation there are a number of potential outcomes of CDK inhibitor treatment, including a harmful increase in reactivation, decreased tumor formation through prevention of entry into latency, or enhanced efficacy of antivirals. Their potential impact on tumor development or established tumors is difficult to predict, but it is possible that increased reactivation may lead to lysis of latently infected tumor cells. This effect, coupled with the decrease in cell division from CDK inhibition itself, may increase efficacy. This potential effect should be considered as therapeutic strategies that increase viral reactivation, coupled with antiviral therapy, have been proposed for the treatment of EBV-associated malignancies.^73^ Given the wide variety of potential outcomes, further model characterization and small animal model studies are needed to direct clinical studies.

In summary, we demonstrated the use of a cell line latently infected with a GFP-reporter virus to identify distinct stages of viral reactivation. We showed that broad spectrum CDK inhibitors consistently inhibit viral production, while specific CDK 4/6 inhibitors make cells more susceptible to reactivation induction. The findings described here provide new insights into viral reactivation as well as the potential clinical applications of CDK inhibitors for the treatment of gammaherpesvirus-associated disease.

## Acknowledgments

We thank the University of Colorado Department of Immunology and Microbiology flow cytometry core and Eric Clambey and members of the van Dyk laboratory for helpful discussions.

